# Dilated Saliency U-Net for White Matter Hyperintensities Segmentation using Irregularity Age Map

**DOI:** 10.1101/550517

**Authors:** Yunhee Jeong, Muhammad Febrian Rachmadi, Maria del C. Valdés-Hernández, Taku Komura

## Abstract

White matter hyperintensities (WMH) appear as regions of abnormally high signal intensity on T2-weighted magnetic resonance image (MRI) sequences. In particular, WMH have been noteworthy in age-related neuroscience for being a crucial biomarker for all types of dementia and brain aging processes. The automatic WMH segmentation is challenging because of their variable intensity range, size and shape. U-Net tackles this problem through the dense prediction and has shown competitive performances not only on WMH segmentation/detection but also on varied image segmentation tasks. However, its network architecture is highly complex. In this study, we propose the use of Saliency U-Net and irregularity age map (IAM) to decrease the U-Net architectural complexity without performance loss. We trained Saliency U-Net using both: a T2-FLAIR MRI sequence and its correspondent IAM. Since IAM guides locating image intensity irregularities, in which WMH are possibly included, in the MRI slice, Saliency U-Net performs better than the original U-Net trained only using T2-FLAIR. The best performance was achieved with fewer parameters and shorter training time. Moreover, the application of dilated convolution enhanced Saliency U-Net by recognising the shape of large WMH more accurately through multi-scale context learning. This network named Dilated Saliency U-Net improved Dice coefficient score to 0.5588 which was the best score among our experimental models, and recorded a relatively good sensitivity of 0.4747 with the shortest training time and the least number of parameters. In conclusion, based on our experimental results, incorporating IAM through Dilated Saliency U-Net resulted an appropriate approach for WMH segmentation.

## 1 INTRODUCTION

White matter hyperintensities (WMH) are commonly identified as signal abnormalities with intensities higher than other normal regions on the T2-FLAIR magnetic resonance imaging (MRI) sequence. WMH have clinical importance in the study and monitoring of Alzheimer’s disease (AD) and dementia progression (Gootjes et al., 2004). Higher volume of WMH has been found in brains of AD patients compared to age-matched controls, and the degree of WMH has been reported more severe for senile onset AD patients than presenile onset AD patients (Scheltens et al., 1992). Furthermore, WMH volume generally increases with the advance of age (Raz et al., 2012; Jagust et al., 2008). Due to their clinical importance, various machine learning approaches have been implemented for the automatic WMH segmentation (Admiraal-Behloul et al., 2005; Bowles et al., 2017).

Limited One-Time Sampling Irregularity Map (LOTS-IM) is an unsupervised algorithm for detecting tissue irregularities, that successfully has been applied for segmenting WMH on brain T2-FLAIR images (Rachmadi et al., 2019). Without any ground-truth segmentation, this algorithm produces a map which describes how much each voxel is irregular compared with an overall area. This map is usually called “irregularity map” (IM) or “irregularity age map” (IAM). The concept of this map was firstly suggested in the field of computer graphics to calculate pixel-wise “age” values indicating how weathered/damaged each pixel is compared to the overall texture pattern of an image (Bellini et al., 2016). Rachmadi et al. then proposed a similar approach to calculate the irregularity level of WMH with respect to the “normal” tissue in T2-FLAIR brain MRI (Rachmadi et al., 2017, 2018b). As WMH highlight irregular intensities on T2-FLAIR MRI slices, IAM can be also used for WMH segmentation. Although performing better than some conventional machine learning algorithms, LOTS-IM still underperforms compared to state-of-the-art deep neural networks. This is mainly because IAM essentially indicates irregular regions, including artefacts, other pathological features and some grey matter regions, in addition to WMH. However, considering IAM depicts irregularities quite accurately and can be generated without a training process, we propose to use IAM as an auxiliary guidance map of WMH location for WMH segmentation.

Recently, the introduction of deep neural networks, the state-of-art machine learning approach, has remarkably increased performances of image segmentation and object detection tasks. Deep neural networks outperform conventional machine learning approaches in bio-medical imaging tasks as well as general image processing. For example, Ciresan et al. built a pixel-wise classification scheme that uses deep neural networks to identify neuronal membranes on electron microscope (EM) images (Ciresan et al., 2012). In another study, Ronneberger et al. proposed a new deep neural network architecture called U-Net for segmenting neuronal structures on EM images (Ronneberger et al., 2015).

In medical images’ segmentation tasks, U-Net architecture and its modified versions have been massively popular due to the end-to-end segmentation architecture and high performance. For instance, a U-Net-based fully convolutional network was proposed to automatically detect and segment brain tumors using multi-modal MRI data (Dong et al., 2017). A 3D U-Net for segmenting the kidney structure in volumetric images produced good quality 3D segmentation results (Ҫiҫek et al., 2016). UResNet, which is a combination of U-Net and a residual network, was proposed to differentiate WMH from stroke lesions (Guerrero et al., 2018). Zhang et al. trained a randomly initialised U-Net for WMH segmentation and improved the segmentation accuracy by post-processing the network’s results (Zhang et al., 2018b).

While there have been many studies showing that U-Net performs well in image segmentation, it has one shortcoming that is long training time due to its high complexity (Briot et al., 2018; Zhang et al., 2018a). To ameliorate this problem, Karargyros et al. suggested the application of regional maps as an additional input, for segmenting anomalies on CT images, and named their architecture Saliency U-Net (Karargyros and Syeda-Mahmood, 2018). They pointed out that extraction of relevant features from images unnecessarily demands very complex deep neural network architectures. Thus, despite neural networks architecture with large number of layers being able to extract more appropriate features from raw image data, it often accompanies a long training time and causes overfitting. Saliency U-Net has regional maps and raw images as inputs, and separately learns features from each data. The additional features from regional maps add spatial information to the Saliency U-Net, which successfully delineates anomalies better than the original U-Net with less number of parameters (Karargyros and Syeda-Mahmood, 2018).

Another way to improve the segmentation performance of deep neural networks is through the recognition of the multi-scale context image information. Multi-scale learning is important particularly for detection/segmentation of objects with variable sizes and shapes. A dilated convolution layer was proposed to make deep neural networks learn multi-scale context better (Yu et al., 2017). Using dilated convolution layers, an architecture can learn larger receptive fields without significant increase in the number of parameters. Previous studies have reported improvements using dilated convolution layers in medical image processing tasks (Lopez and Ventura, 2017; Moeskops et al., 2017).

In this paper, we propose to use IAM as an additional input data to train a U-Net neural network architecture for WMH segmentation, owed to the fact that LOTS-IM can easily produce IAM without the need for training using manually marked WMH ground-truth data. U-Net architecture is selected as a base model for our experiments as it has shown the best learning performance using IAM (Rachmadi et al., 2018a). To address the incorporation of IAM to U-Net for WMH segmentation, we propose feed-forwarding IAM as regional map to a Saliency U-Net architecture. We also propose combining Saliency U-Net with dilated convolution to learn multi-scale context from both T2-FLAIR MRI and IAM data, in a scheme we name Dilated Saliency U-Net. We compare the original U-Net’s performance with the performances of Saliency U-Net and Dilated Saliency U-Net on WMH segmentation.

Consequently, the contributions of our work can be summarised as follows:

- Proposing the use of IAM as an auxiliary input for WMH segmentation. T2-FLAIR MRI and IAM complement each other when they both are used as input to the neural network, addressing challenging cases especially those with few small WMH.
- Integration of Saliency U-Net and dilated convolution for WMH segmentation; which showed more detailed boundary delineation of large WMH. It also attained the best Dice coefficient score compared to our other experimental models.

## 2 MATERIALS AND METHODS

### 2.1 Dataset

MRI can produce different types of images to display normal tissues and different types of clinical abnormalities. It is desirable to choose suitable image types considering the properties of biomarkers or diseases targeted in the segmentation task. T2-weighted is one of the MRI sequences that emphasises fluids as bright intensities. The bright intensity of fluids makes WMH difficult to identify in this MRI modality because WMH are also bright on T2-weighted. T2-fluid attenuated inversion recovery (T2-FLAIR) removes cerebrospinal fluid (CSF) signal from the T2-weighted sequence, increasing the contrast between WMH and other brain tissues. Therefore, we have chosen T2-FLAIR MRI as the main source of image data for our experiments.

We obtained T2-FLAIR MRI sequences from the public dataset *the Alzheimer’s Disease Neuroimaging Initiative* (ADNI)^1^ which was initially launched by Mueller et al. (Mueller et al., 2005). This study has mainly aimed to examine combinations of biomarkers, MRI sequences, positron emission tomography (PET) and clinical-neuropsychological assessments in order to diagnose the progression of mild cognitive impairment (MCI) and early AD. From the whole ADNI database, we randomly selected 60 MRI scans collected for three consecutive years from 20 subjects with different degrees of cognitive impairment in order to evaluate the applicability of our proposed scheme not only for cross-sectional studies but also for longitudinal analyses of WMH. Each MRI scan has dimensions of 256 × 256 × 35. We describe how train and test dataset are composed in Section 2.8.

Ground truth masks were semi-automatically produced by an experienced image analyst using a thresholding algorithm combined with region-growing in the Object Extractor tool of Analyze™ software. This semi-automatic WMH segmentation used the T2-FLAIR images. Intracranial volume (ICV) and CSF masks were generated automatically using optiBET (Lutkenhoff et al., 2014), and a multispectral algorithm developed in-house (Hernández et al., 2015) respectively. Full details and binary WMH reference masks can be downloaded from the University of Edinburgh DataShare repository^2^.

### 2.2 Irregularity Age Map (IAM)

As described in Section 1, the concept of IAM was proposed with the development of the LOTS-IM algorithm and its application to the task of WMH segmentation (Rachmadi et al., 2017, 2018b, 2019). This algorithm was inspired by the concept of “age map” proposed by Bellini and colleagues while calculating the level of weathering or damage of pixels compared to the overall texture pattern on natural images (Bellini et al., 2016). Rachmadi et al. adopted this principle to compute the degree of irregularity in brain tissue from T2-FLAIR MRI.

In this study, the GPU-powered LOTS-IM algorithm (Rachmadi et al., 2019)^3^ was used to generate IAM from all scans. The steps of the LOTS-IM algorithm are as follows. Source and target patches are extracted from the MRI slices with four different sizes (i.e., 1 × 1, 2 × 2, 4 × 4 and 8 × 8) to capture different details in the brain tissues (Rachmadi et al., 2017). All grid fragments consisting of *n × n* sized patches are regarded as *source patches*. On the other hand, *target patches* are picked at random locations within the brain. Thus, non-brain target patches, located within the CSF mask or outside the ICV mask, are excluded from computation. Then, the difference between each source patch and one target patch on the same slice is calculated by Eq 1;

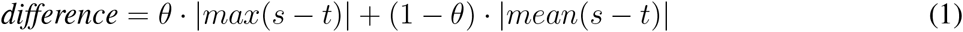

where *s* and *t* mean source patch and target patch respectively, also *θ* was set to 0.5 (Rachmadi et al., 2018b). After difference values between a source patch and all target patches are calculated, the 100 largest difference values are averaged to become the *age value* of the corresponding source patch (Rachmadi et al., 2017). The rationale is that the average of the 100 largest difference values produced by an “irregular” source patch is still comparably higher than the one produced by a “normal” source patch (Rachmadi et al., 2017, 2018b). Furthermore, the age value is computed only for source patches within the brain to reduce the computational complexity. All age maps from four different patch sizes are, then, normalised to have normalised age values between 0 and 1; and each of them is up-sampled into its original image size and smoothed by a Gaussian filter. The final age map is produced by blending these four age maps using the Eq 2;

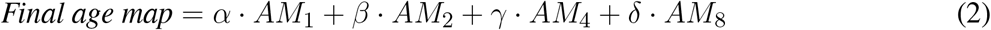

where *AM*_*x*_ means the age map of *x* × *x* sized patches and *α* + *β* + *γ* + *δ* = 1. In this study, *α* = 0.65, *β* = 0.2, *γ* = 0.1 and *δ* = 0.05 (Rachmadi et al., 2019). Finally, the final age map is penalised by multiplying the original T2-FLAIR image slice to reflect only the high intensities of WMH, and globally normalised from 0 to 1 over all brain slices. The overall steps are schematically illustrated in Figure 1.

**Figure 1.**
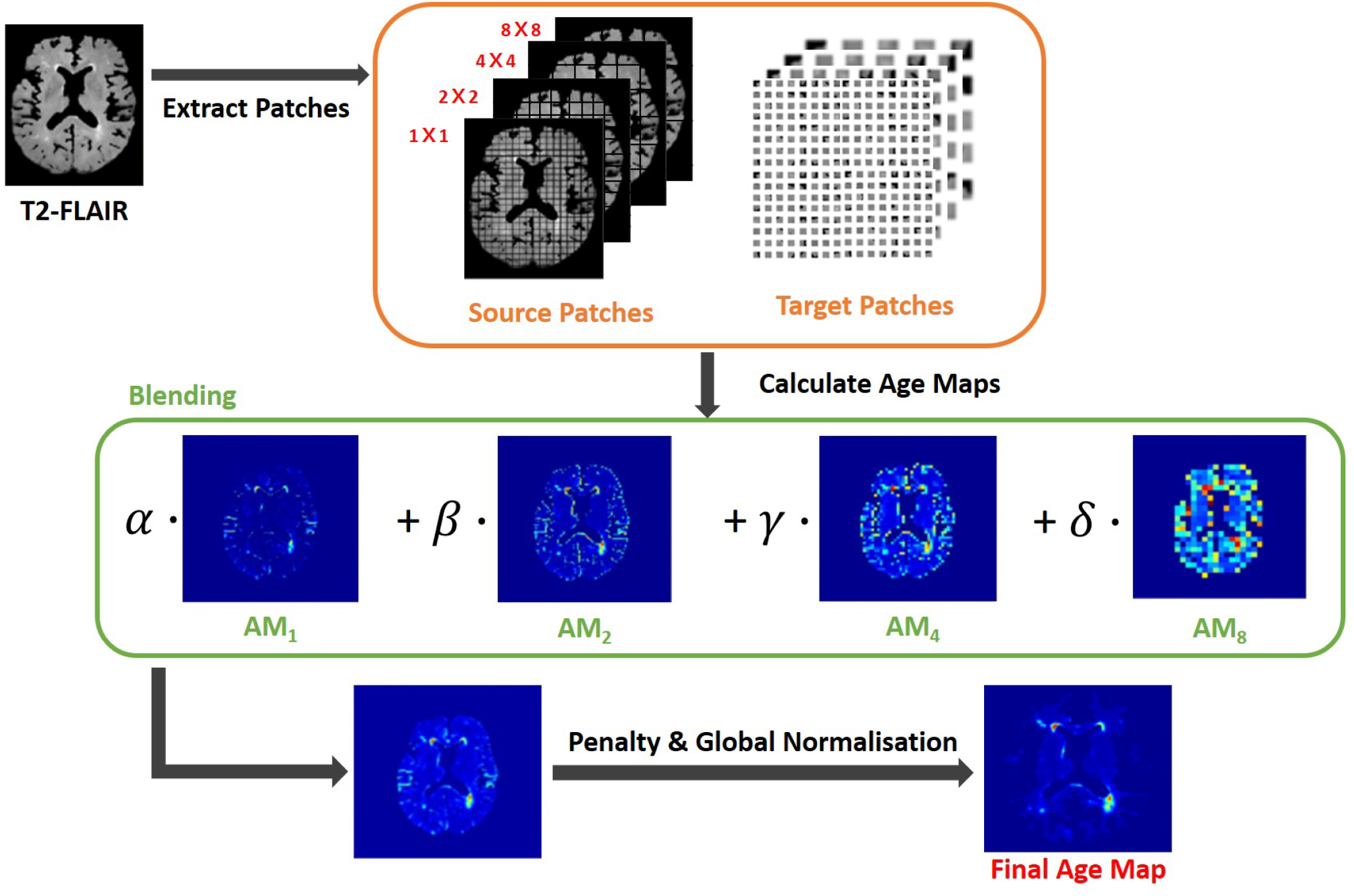
Flow chart illustrating the LOTS-IM algorithm proposed by Rachmadi and colleagues (Rachmadi et al., 2019) applied to WMH segmentation. This study uses the final map generated by this algorithm, and refers to it as “IAM data”.

Though regarded as WMH segmentation map in the original studies, IAM essentially calculates the probability of each voxel to constitute an irregularity of the “normal” tissue. This irregular pattern includes not only WMH but more features such as artefacts, T2-FLAIR hyperintensities of other nature, as well as sections of the cortex that could be hyperintense. To compensate these flaws and take advantage of its usefulness, we developed a new scheme that uses IAM as an auxiliary guidance map for training deep neural networks rather than using it for producing the final WMH segmentation.

### 2.3 U-Net

Since U-Net architecture was firstly presented (Ronneberger et al., 2015), various image segmentation studies have used this architecture due to its competitive performance regardless of the targeted object types. Different to the natural image segmentation, bio-medical image segmentation involves a more challenging circumstance as lack of data for the training process is a common problem. U-Net deals with this challenge with dense prediction of the input image using up-sampling layers that produce equal-sized input and output. This approach was drew by fully convolutional networks (Long et al., 2015).

U-Net is comprised of two parts, the encoding part where feature maps are down-sampled by max-pooling layers and the decoding part where the reduced size of feature maps are up-sampled to the original size. It retains the localisation accuracy with the contracting path, which concatenates the feature maps stored in the encoding part with the decoding part. These kept high resolution features help to restore the details of localisation removed by max-pooling layer, when the feature maps are up-sampled in the decoding part. The architecture is depicted in Figure 3 (a).

A drawback of U-Net is its large number of parameters. To restore the high resolution localisation, the network should increase the number of feature channels in the decoding part. Training time and memory usage are proportional to the number of parameters. So training a U-Net architecture is constrained by its high consumption of time and memory. Moreover, the complexity of the (neural) network often induces the problem of overfitting.

### 2.4 Saliency U-Net

Saliency U-Net was first introduced to detect anomalies in medical images using a combination of raw (medical) images and simple regional maps (Karargyros and Syeda-Mahmood, 2018). Saliency U-Net performed better than U-Net while using less number of parameters. An architecture with less number of parameters is preferable as it is easier and faster to be trained. Karargyros and Syeda-Mahmood showed that convolution layers are not needed to extract more relevant features from raw images if auxiliary information from regional map is given as input. The Saliency U-Net architecture has two branches of layers in the encoding part (Figure 3 (b)). Each branch extracts features from raw image and regional map independently, and the extracted features are fused before the decoding part.

Segmentation results from Saliency U-Net in the original study (Karargyros and Syeda-Mahmood, 2018) showed more precise localisation and better performance than the original U-Net, which contained a larger number of convolutional layers. Therefore, for WMH segmentation, we propose to use Saliency U-Net taking T2-FLAIR as raw input image and IAM as regional map.

### 2.5 Dilated Convolution

One common issue for image segmentation via deep neural networks is caused by the reduced size of the feature maps in the pooling layer introduced to capture global contextual information. While pooling layers are useful to get rid of some redundancies in feature maps, the lower size of feature maps after the last pooling layer also causes loses of some of its original details/information, decreasing the segmentation performance where the targeted regions are not spatially prevalent(Hamaguchi et al., 2018; Yu et al., 2017).

Dilated convolution solved this problem by calculating a convolution over a larger region without reducing the resolution (Yu and Koltun, 2015). The dilated convolution layer enlarges a receptive field including *k* skips between each input pixel. *k* is called *dilation factor*. In numerical form, a dilated convolution layer with a dilation factor *k* and a *n* × *n* filter is formulated as follows:

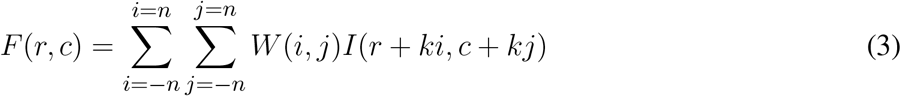

Figure 2 (a)-(c) show examples of dilated convolution filters with dilation factors 1 to 3.

**Figure 2.**
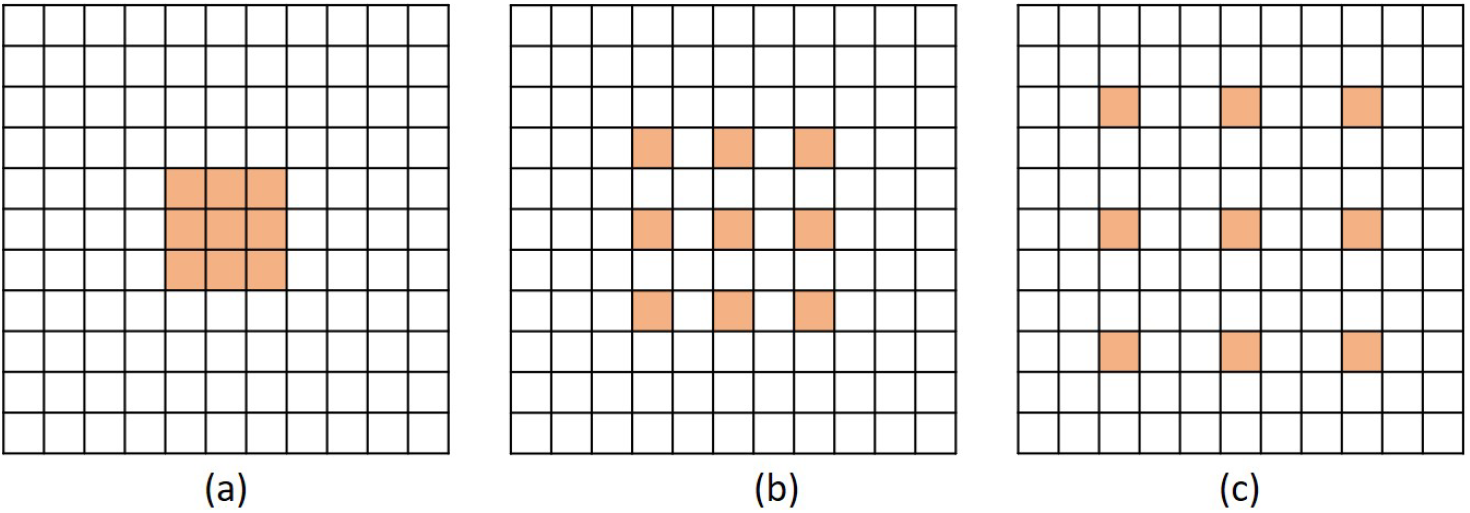
Examples of dilated convolution filter with 3 × 3 size. (a) Dilation factor = 1, (b) Dilation factor = 2 and (c) Dilation factor = 3.

The additional advantage of dilated convolution is to widen the receptive field without increasing the number of parameters. Large receptive fields learn the global context by covering a wider area over the input feature map, but bring a memory leak and time consumption out for a growing number of parameters. Dilation can expand the receptive field of the convolution layer as much as skipped pixels without extra parameters. For instance, as shown in Figure 2 (a) and (c), the filter with dilation factor 3 has 7 × 7 sized receptive field, while the filter with dilation factor 1 has 3 × 3 sized receptive field.

In this study, we propose the incorporation of dilated convolution to Saliency U-Net for WMH segmentation. Since the size of WMH is variable, it is necessary to recognise different sizes of spatial contexts for more accurate delineation of WMH. We believe that dilated convolutions can manage the variable size of WMH from different sizes of receptive field.

### 2.6 Our Experimental Models

We examined three different U-Net models for which its original architecture was trained using input data with different modalities: T2-FLAIR (model 1), IAM (model 2) and both (model 3). To feed both T2-FLAIR and IAM together, we integrated T2-FLAIR and IAM as a two-channel input. As mentioned in Section 2.3, U-Net architecture has encoding and decoding parts. In the encoding part, input images or feature maps are down-sampled by max-pooling layers to obtain relevant features for WMH segmentation. Then, in the decoding part, reduced feature maps are up-sampled again by up-sampling layers to acquire the original size in the final segmentation map. Max-pooling and Up-sampling layers are followed by two CONV blocks (yellow blocks in Figure 3). The CONV block contains a convolution layer, an activation layer and a batch normalisation layer. Batch normalisation allows to train neural networks with less careful initialisation and higher learning rate by performing normalisation at every batch (Ioffe and Szegedy, 2015). All activation layers except the last one are ReLU (Nair and Hinton, 2010), but the last activation layer calculates the categorical cross-entropy to yield a probability map for each label.

**Figure 3.**
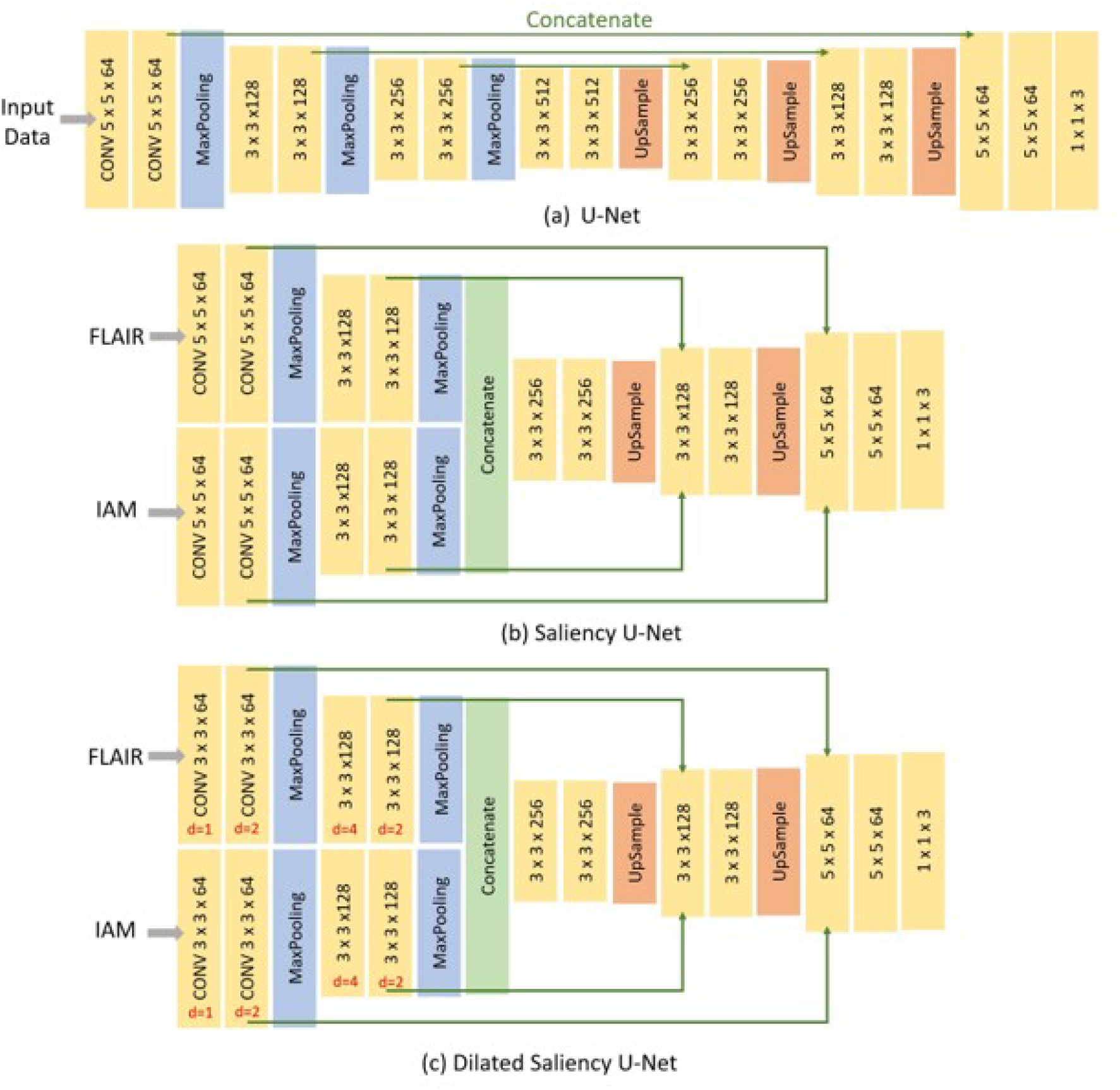
Architecture of three different networks used in this study. (a) the original U-Net, (b) Saliency U-Net and (c) Dilated Saliency U-Net. Three numbers of CONV block (yellow block) represents *filter size* × *filter size* × *filter channels*. For the Dilated Saliency U-Net model, red numbers mean a dilation factor for the convolution layer in each CONV block.

In addition, we trained Saliency U-Net and Dilated Saliency U-Net by feed forwarding both T2-FLAIR and IAM separately. In this way, we assume that IAM works as a simple regional map which provides localisation information of WMH rather than just being a different image channel. While the U-Net architecture has one branch of the encoding part, Saliency U-Net encoding part consists of two branches that learn raw images and regional maps individually. Furthermore, we applied dilation factors of 1, 2, 4 and 2 to the first four convolutional layers of Saliency U-Net to form the Dilated Saliency U-Net. The architectures of U-Net, Saliency U-Net and Dilated Saliency U-Net can be seen in Figure 3.

Performance of these models are compared to each other in Section 3. We additionally conducted experiments on the original U-Net models trained only with T2-FLAIR and only with IAM in order to see how using both T2-FLAIR and IAM as inputs affects learning WMH segmentation. Our five experimental models are listed in Table 1.

**Table 1.** Dice Similarity Coefficient (DSC), sensitivity, positive predictive value (PPV), training time and number of parameters for our five experimental models. Values in bold are the highest scores and in italic the second highest. In the brackets after the model names, the input data type is specified. “FLAIR” is equivalent to T2-FLAIR and “F+I” refers to taking both T2-FLAIR and IAM as input.

| Model | DSC | Sensitivity | PPV | Training Time | # Parameters |
| --- | --- | --- | --- | --- | --- |
| U-Net(FLAIR) | 0.5440 | 0.4594 | 0.6275 | 1h 52m 55s | 7,859,715 |
| U-Net(IAM) | 0.5274 | 0.4179 | **0.6769** | 1h 53m 52s | 7,859,715 |
| U-Net(F+I) | 0.5281 | **0.4902** | 0.6268 | 1h 24m 22s | 7,861,315 |
| Saliency U-Net(F+I) | *0.5535* | 0.4730 | 0.6034 | 1h 30m 1s | 2,756,803 |
| Dilated Saliency U-Net(F+I) | **0.5588** | *0.4747* | *0.6374* | 1h 4m 18s | 2,623,683 |

### 2.7 Preprocessing

In machine learning, data preprocessing is needed to standardise the data into a comparable range. It is especially important when we deal with MRI data whose intensity is not in a fixed range. Differences in the intensity range are caused by differences in MRI acquisition protocols, scanner models, calibration settings, etc. (Shah et al., 2011).

For this reason, we normalised the intensity of the brain tissue voxels in our train and test data. The image intensity of the majority of non-brain tissue voxels of an MRI slice is zero or near-zero, although few non-brain voxels can have peak intensity values above the intensity range of the brain tissue. Thus normalising intensities from all voxels together can bias the intensity values towards zero and reduce the effect of WMH on brain tissue voxels. Brain tissue voxels were filtered using CSF and the intracranial volume (ICV) masks as follows:

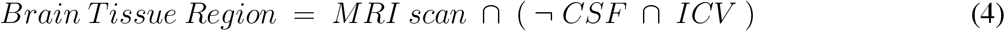

We normalised the brain tissue voxels on each slice into a distribution with zero-mean and unit variance by subtracting the mean value from each voxel value and dividing the result by the standard deviation.

Although WMH segmentation can be regarded as the binary classification of voxels, we re-labelled the ground-truth data assigning voxels one of the three following labels: non-brain, non-WMH brain tissue and WMH. However, when evaluating the segmentation results, we considered both non-brain and non-WMH brain tissue labels as non-WMH labels to calculate sensitivity and Dice similarity coefficient which are metrics for the binary classification. Figure 4 shows the example of a T2-FLAIR slice, the same slice after preprocessing and normalisation, and the ground-truth slice.

**Figure 4.**
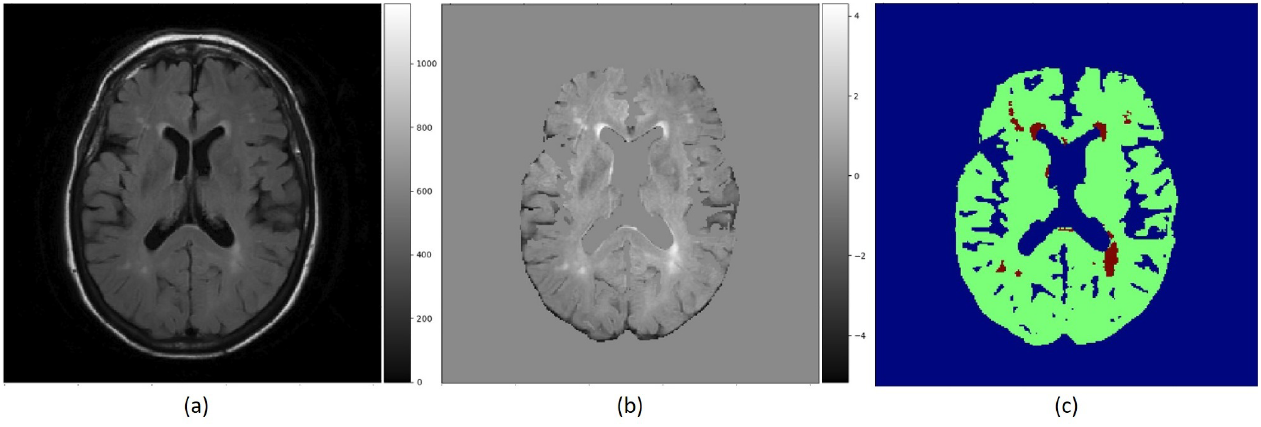
(a) Raw T2-FLAIR image, (b) T2-FLAIR input after preprocessing and normalisation, (c) Ground truth data with three labels. Blue region is non-brain area, green region is non-WMH brain tissues and red region is WMH.

### 2.8 Training and Testing Setup

For training, 30 MRI scans of the ADNI dataset described in Section 2.1 were randomly selected. These 30 MRI scans were collected from 10 subjects for three consecutive years. We trained our networks with image patches generated from these MRI scans, not slices, to increase the amount of training data. If we train our models using slice images, the amount of training data is only 35 × 30 = 1050 slices, which is not ideal for training a deep neural network architecture. Instead, by extracting 64 × 64 sized patches from each image slice, we could have 30,000 patches for training data.

For testing, we used the rest 30 scans of the ADNI sample, which are not used during training. These scans were also obtained from another 10 subjects for three consecutive years. The testing dataset was comprised of image slices without patch extraction. Slice image data is necessary to analyse the results from our models according to the distributions or volumes of WMH. Our testing dataset holds 1050 of 256 × 256 image slices in total as each scan contains 35 slices.

All experimental models were trained using the same network configuration. We set learning rate to 1*e*^−5^ and batch size to 16. As an optimisation method, we selected the Adam optimisation algorithm (Kingma and Ba, 2014), although the original U-Net scheme used the stochastic gradient descent (SGD) optimiser. This is because the Adam optimiser can handle sparse gradients. It is highly possible that our training data produce sparse gradients as non-brain voxels, which are the majority, have zero intensity. We applied the Adam optimiser accordingly, considering this data property.

## 3 RESULTS

In this section, we present how experiments were conducted, and analyse and compare the experimental results.

### 3.1 Evaluation Metrics

We use sensitivity, positive predictive value (PPV) and Dice similarity coefficient (DSC) to evaluate the models. Sensitivity measures the rate of true positives as below:

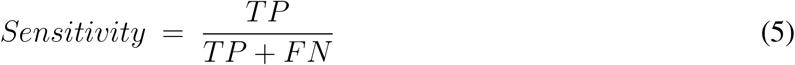

where *TP* means true positive, and *FN* means false negative. PPV also measures the rate of true positives but from the total of positive calls like below:

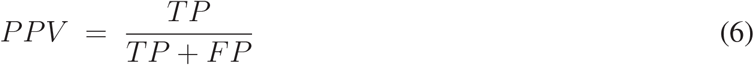

where FP refers to false positive. DSC is a statistic method to compare the similarity between two samples of discrete values (Dice, 1945). It is one of the most common evaluation metrics in image segmentation.

The formula is as follow:

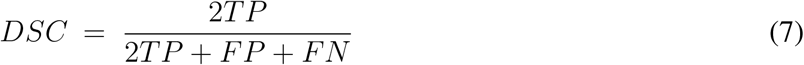

where *TP* and *FN* are as per Equation 5 and *FP* means false positive. DSC is interpreted as the overlapping ratio to the whole area of prediction and target objects, while sensitivity measures the correctly predicted region of the target object. If the prediction includes not only true positives but also wrong segmentation results (false positives), the DSC score can be low despite the high sensitivity.

### 3.2 The Effects of IAM as an Auxiliary Input Data

Table 1 shows overall performances of our five experimental models. The adoption of IAM as an auxiliary input data for U-Net (i.e., U-Net(F+I)) improved sensitivity to 0.4902 but had lower DSC score than the model that used only the T2-FLAIR image as input. On the other hand, Saliency U-Net(F+I) improved the DSC scores achieved by U-Net to 0.5535 while Dilated Saliency U-Net(F+I) achieved the best DSC score of 0.5588. Dilated Saliency U-Net(F+I) yielded the second best sensitivity rate after U-Net trained with T2-FLAIR and IAM (i.e., U-Net(F+I)). U-Net(IAM) achieved the best PPV value of our five models and Dilated Saliency U-Net(F+I) achieved the second highest value of PPV. From these results, we can see that the three models trained with T2-FLAIR and IAM particularly increased the sensitivity performance of the network architectures.

Saliency and Dilated Saliency U-Net included considerably less parameters than the three U-Net models. As shown in Table 1, Saliency and Dilated Saliency U-Net have more than three times less parameters and slightly shorter training time than the original U-Net while having better if not similar performance on WMH segmentation.

With regards to training time, although feeding both T2-FLAIR and IAM together into U-Net involved the calculation of more parameters due to the two-channel input, the training time for this model was shorter than that of U-Net(FLAIR) and U-Net(IAM). In deep learning studies, visual attention, which gives larger weight on the region of interest, speeds up learning by leading the model to concentrate on the relevant regions. This has been experimentally demonstrated in previous studies (Najibi et al., 2018; Choi et al., 2017). In our case, IAM confers the visual attention effect to the network architecture. Despite having fewer parameters, Saliency U-Net took longer time to train than U-Net(F+I). Feed-forward and back-propagation proceed separately in each encoding part. Dilated Saliency U-Net significantly decreased the training time compared to the other models by skipping voxels that reduce the computational complexity, when calculating the convolution.

Figure 5 presents training and validation losses for our five models. Same colour lines correspond to the same model. Solid and dashed lines represent training loss and validation loss each. For all models, both training and validation losses properly converged. Thus, our models are not overfitted on the training data.

**Figure 5.**
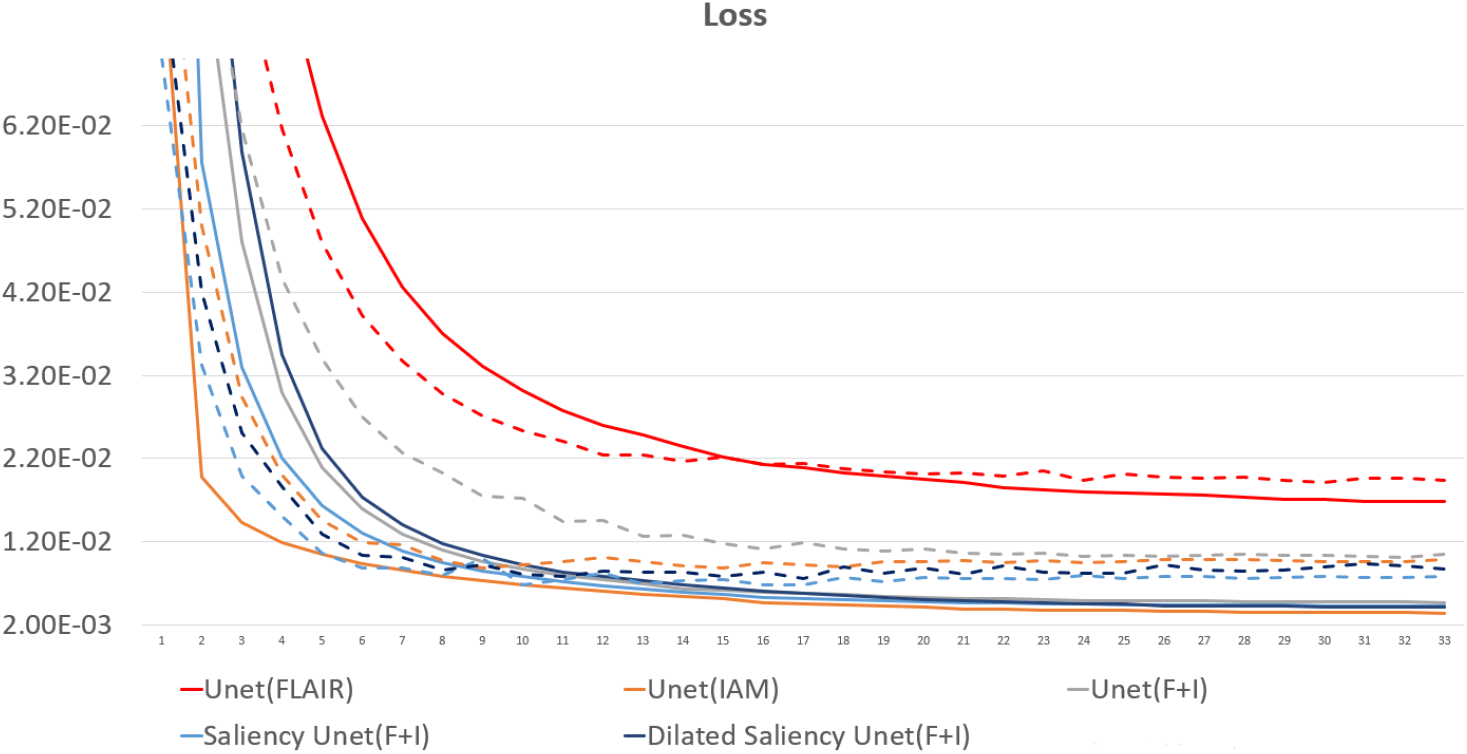
Loss graph of our five models. While solid lines indicate training loss, dashed lines represent validation loss.

We also evaluated whether the median and the distribution of DSC scores throughout the testing set differed significantly between the five models evaluated. We conducted two tests: 1) the Wilcoxon ranksum, as implemented by the function `ranksum` in MATLAB, to evaluate whether the medians of the DSC scores from each model across the testing dataset were significantly different between each other; and 2) the Kruskal-Wallis test, as implemented by the MATLAB function `kruskalwallis`, to evaluate whether the distributions of these DSC values were statistically significantly different between the models. Neither the medians nor the DSC distributions obtained by these five models significantly differed. The result of the Kruskal-Wallis test is shown in Table 2. The *p-value* obtained from the ANalysis Of VAriance (ANOVA) of the DSC distributions from the five models across all cases is 0.7786, indicating that the results of these five models did not differ significantly from each other in terms of the distribution of DSC across the testing set. This emphasises that Dilated Saliency U-Net model can produce similar level of performance as the original U-Net models even with less number of parameters and shorter training time. Figure 6 also illustrates that the DSC scores obtained from applying our models are similarly distributed to each other.

**Table 2.** ANOVA table for our five models. SS refers to the sum of squares. df and MS mean degrees of freedom and mean squares respectively.

| Source | SS | df | MS | $F$-value | $p$-value |
| --- | --- | --- | --- | --- | --- |
| Models | 3334.7 | 4 | 833.68 | 1.77 | 0.7786 |

**Figure 6.**
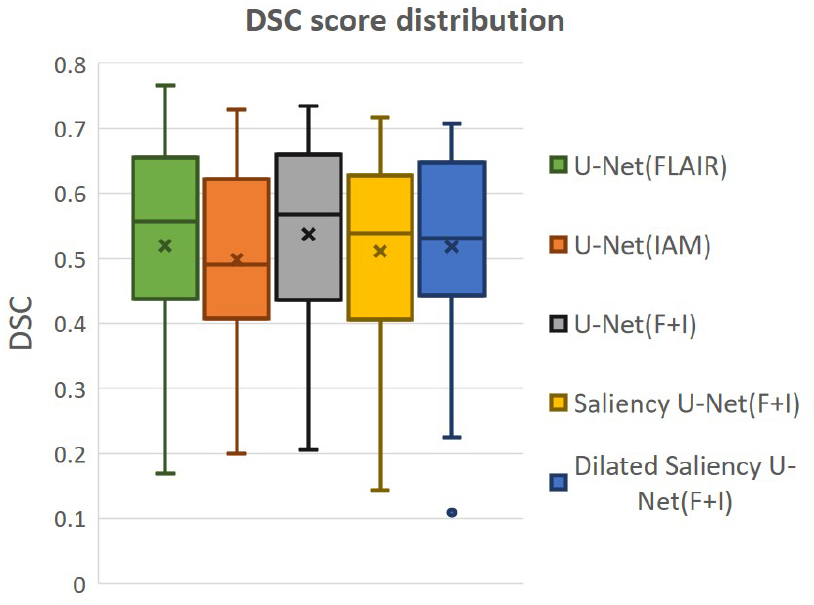
Distributions of DSC score by our five models.

Figure 7 visualises the examples of WMH segmentation results by our experimental models. In most cases, the use of two data sources (i.e., IAM and T2-FLAIR images) in training the network complements each other’s effect detecting tricky WMH regions. Depending on the contrast/size of WMH or the quality of IAM, there are some cases in which WMH are distinguishable on IAM but unclear in T2-FLAIR and vice versa. For example, if WMH clusters are too small, it is hard to differentiate them on T2-FLAIR, but they are better observable on IAM, where WMH and normal brain tissue regions have better contrast. On the other hand, in the presence of other irregular patterns such as extremely low intensities of brain irregularities around WMH, T2-FLAIR can indicate WMH clearly than IAM. In Figure 7 Row (a), U-Net(FLAIR) produced better WMH segmentation result than U-Net(IAM) due to the poor quality of IAM. Conversely, U-Net(FLAIR) could not detect WMH well due to unclear intensity contrast on T2-FLAIR while U-Net(IAM) could segment these WMH regions as IAM enhanced them as anomalies (Figure 7 Row (b)). Furthermore, incorporating both T2-FLAIR and IAM together as input data produced better WMH segmentation in general (5^*th*^-7^*th*^ columns from left to right of Figure 7).

**Figure 7.**
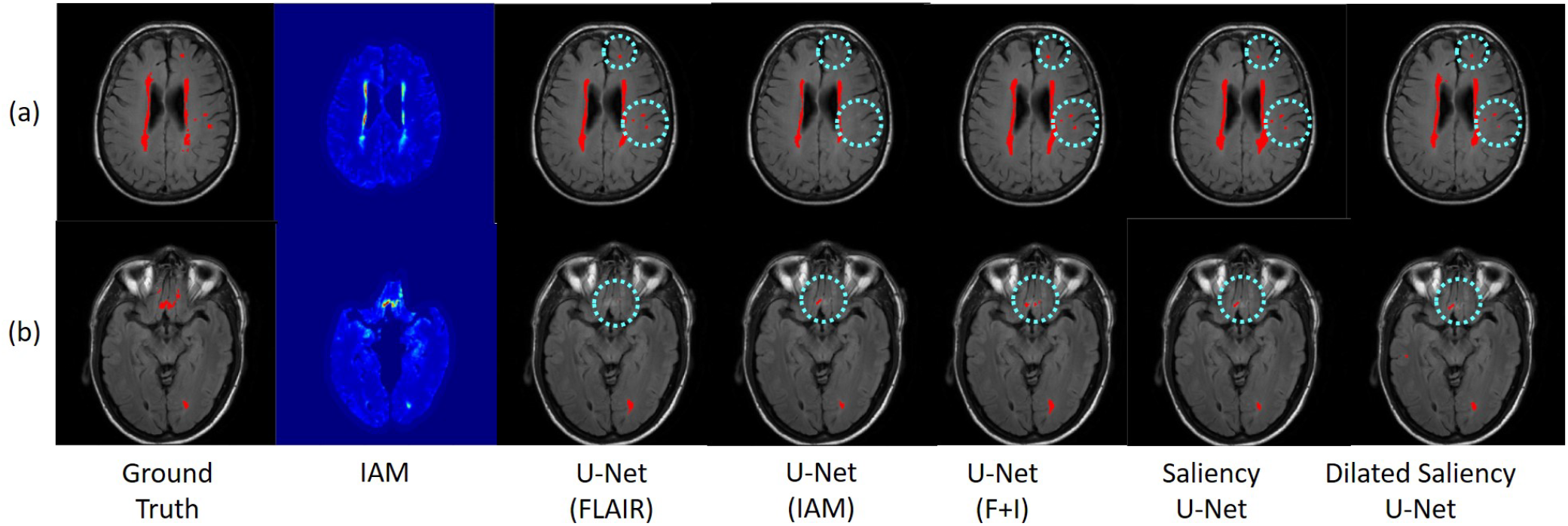
Examples of WMH segmentation results by our experimental models. Cyan circles indicate WMH detected only by one of the original U-Net models, i.e. U-Net(FLAIR) or U-Net(IAM). Row (a) shows a case where WMH is distinguishable exclusively in the T2-FLAIR image, while row (b) shows a case where IAM highlights WMH clearly. By training networks using T2-FLAIR and IAM, both WMH regions are detected.

### 3.3 WMH Volume analysis

In this experiment, we evaluate our models based on the WMH volumes of the MRI scan (i.e., WMH burden) to examine the influence of WMH burden on the performance of WMH segmentation. The WMH volume of each MRI scan is calculated by multiplying the number of WMH voxels by the voxel size. We grouped MRI scans into three groups according to the range of WMH volume. Table 3 shows the range of WMH volume used as criteria for forming the groups, and the number of scans included in each group. Figure 8 (a) shows the lack of ambiguity or overlap in the classification of the MRI scans in each group.

**Table 3.**
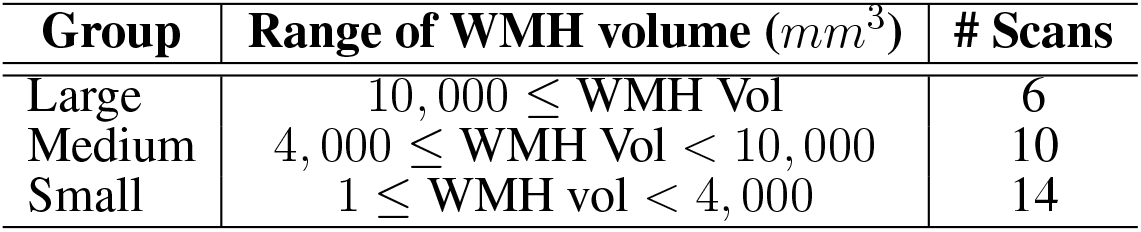
Criteria sorting MRI scans according to WMH voxel volume. “# Scans” means the number of included MRI scans. Most of scans are included in Small and Medium groups.

| Group | Range of WMH volume ($mm^3$) | # Scans |
| --- | --- | --- |
| Large | $10,000 \leq \text{WMH Vol}$ | 6 |
| Medium | $4,000 \leq \text{WMH Vol} < 10,000$ | 10 |
| Small | $1 \leq \text{WMH vol} < 4,000$ | 14 |

**Figure 8.**
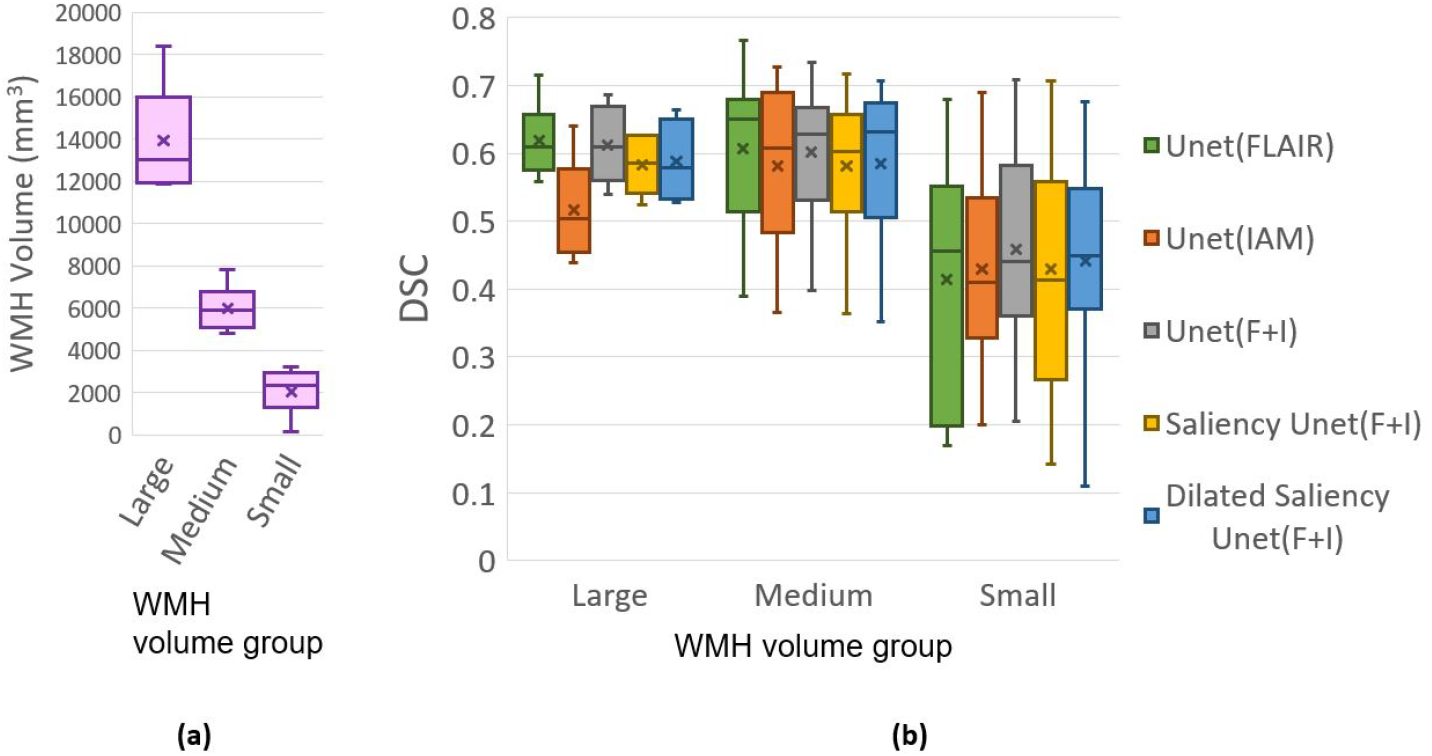
(a) Distributions of data (MRI scan) grouped together based on WMH volume. (b) DSC distributions yielded by tested five models based on WMH volume. “x” and bar at the middle of box indicate mean and median each. Bottom and top of each box mean the first and third quartile.

Figure 8 (b) plots the DSC scores yielded by the MRI scans in the different WMH volume groups by our five experimental models. Please, note that the DSC scores referred in this section correspond to the evaluation of the WMH segmentation results in each MRI scan, not per slice which are used for overall performance evaluation in Section 3.2 Table 1. Hence scans of the Large group might have several small WMH rather than one large region with confluent WMH.

All models tested in this study showed high median values of DSC scores in the Medium group, for which all models performed better than the other groups. In the Large group, U-Net(FLAIR) and U-Net(F+I) models performed similarly well, while U-Net(IAM) performed worst compared with the rest of the models. Mean, median and standard deviation (std.) values of DSC score distribution in each group are shown in Table 4. Overall, the performance of the models for scans with Small and Medium WMH burden was quite similar (see also Figure 8 (b)). However, large variations in DSC scores were observed among the scans of the Small group, especially for the U-Net(FLAIR) model.

**Table 4.** Mean, median and standard deviation values of the distributions of DSC scores from our experimental models per WMH volume groups. Model name *DSU-Net abcd* refers to Dilated Saliency U-Net model with dilation factors *a*, *b*, *c*, *d* in order from the first to the fourth convolution layers, and its trend of dilation factor pattern is specified in the bracket. These dilation factors are applied on convolution layers in the encoding part (i.e., before concatenating T2-FLAIR and IAM feature maps) of the CONV blocks, which consists of convolution, ReLU, and batch normalisation layers. These different Dilated Saliency U-Net models are described in Section 3.6. DSU-Net 1242 was used for the Dilated Saliency U-Net model evaluated in Section 3.3.

| Model | DSC Mean. |  |  | DSC Median. |  |  | DSC std. |  |  |
| --- | --- | --- | --- | --- | --- | --- | --- | --- | --- |
|  | Large | Medium | Small | Large | Medium | Small | Large | Medium | Small |
| U-Net(FLAIR) | 0.6184 | 0.6070 | 0.4147 | 0.6987 | 0.6499 | 0.4559 | 0.0524 | 0.1076 | 0.1746 |
| U-Net(IAM) | 0.5168 | 0.5817 | 0.4294 | 0.5036 | 0.6080 | 0.4106 | 0.0668 | 0.1111 | 0.1455 |
| U-Net(F+I) | 0.6124 | 0.6025 | 0.4580 | 0.6092 | 0.6276 | 0.4400 | 0.0548 | 0.0931 | 0.1460 |
| Saliency U-Net(F+I) | 0.5824 | 0.5812 | 0.4299 | 0.5853 | 0.6023 | 0.4134 | 0.0377 | 0.0956 | 0.1687 |
| DSU-Net_1224 | 0.5722 | 0.5929 | 0.4003 | 0.5814 | 0.6286 | 0.3876 | 0.0592 | 0.0965 | 0.1733 |
| DSU-Net_4221 | 0.5711 | 0.5768 | 0.4253 | 0.5776 | 0.6152 | 0.4250 | 0.0574 | 0.1097 | 0.1640 |
| DSU-Net_1242 | 0.5882 | 0.5852 | 0.4407 | 0.5782 | 0.6320 | 0.4498 | 0.0536 | 0.1069 | 0.1558 |

### 3.4 Longitudinal Evaluation

In the longitudinal evaluation test, we addressed the capacity of our five models in predicting WMH in subsequent years after being trained only using the first year samples. Hence, the training set was formed by the first year samples while the testing set was composed by the second and third year samples. Table 5 shows the mean DSC score for each sample. In this evaluation, U-Net(IAM) and Saliency U-Net performed slightly better than the other three models, partly owed to IAM which could provide information to predict WMH occurrence. As expected, all our models predicted better WMH in the second year than in the third year.

**Table 5.** DSC score for longitudinal evaluation of our five models. We evaluated these models using data from both second and third years. As per Table 1, values in bold are the highest scores and in italics are the second highest ones.

| Model | 2nd year | 3rd year |
| --- | --- | --- |
| U-Net(FLAIR) | 0.6136 | 0.5878 |
| U-Net(IAM) | **0.6270** | *0.6110* |
| U-Net(F+I) | 0.6229 | 0.5823 |
| Saliency U-Net | *0.6258* | **0.6119** |
| Dilated Saliency U-Net | 0.6060 | 0.5881 |

### 3.5 U-Net vs. Saliency U-Net

In order to evaluate the effectiveness of the Saliency U-Net architecture, we compared the original U-Net and Saliency U-Net models trained with T2-FLAIR and IAM. As shown in Table 1, Saliency U-Net yielded higher DSC score than U-Net(F+I) despite U-Net(F+I) having higher sensitivity value. Figure 9 shows that Saliency U-Net successfully eliminates some of the false positives observed in the segmentation result from U-Net(F+I).

**Figure 9.**
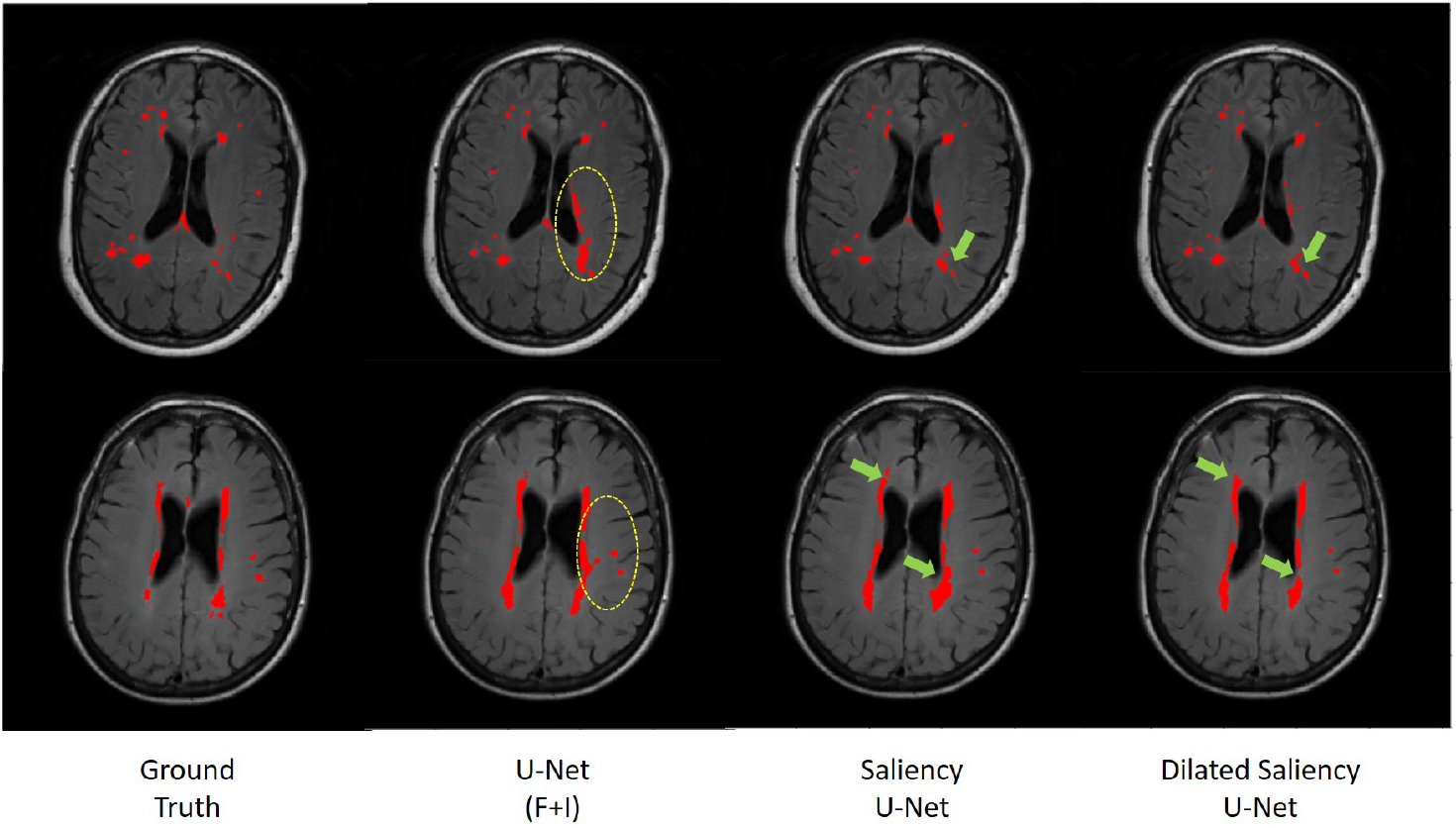
Comparison of WMH segmentation results from U-Net(F+I), Saliency U-Net and Dilated Saliency U-Net. Yellow circles indicate false positive results by U-Net(F+I). These false positive results are eliminated in the results from Saliency and Dilated Saliency U-Net. Green arrows are pointing to locations where boundaries are segmented in more detail by Dilated Saliency U-Net.

We also investigated the change in Saliency U-Net’s performance in relation to its complexity when the number of convolution layers increased/decreased. DSC score, training time and model complexity (i.e., the number of parameters) are compared in Figure 10. The rule for changing the Saliency U-Net complexity is to connect/disconnect the 2 CONV blocks that are attached/detached at both ends, through a “skip” connection. However, since the encoder part is a two-branch architecture, 6 CONV blocks are included at once increasing its complexity (i.e., 4 CONV blocks are added to the encoder part and 2 CONV block are added to the decoder part). Similar approach is done when decreasing the complexity, where 4 CONV blocks and 2 CONV blocks are dropped from the encoder and decoder respectively. For clarity, our original Saliency U-Net model (i.e., evaluated in Table 1 of Section 3.2) contains 14 CONV blocks and each CONV block holds one convolution layer as shown in Figure 3.

**Figure 10.**
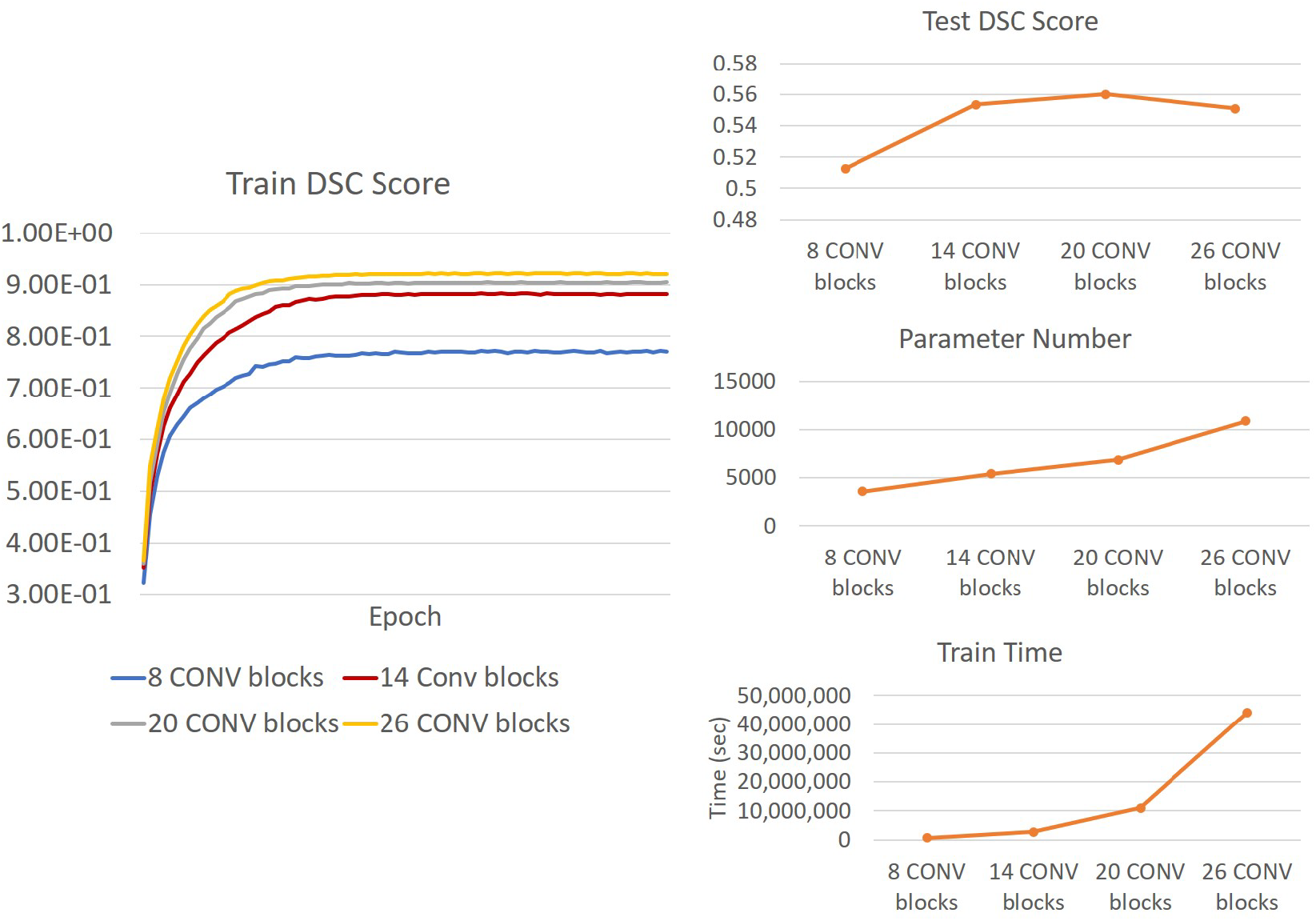
**(Right)** Trends of DSC score, training time and number of parameters of Saliency U-Net when more convolution layers are changed. It is also shown Saliency U-Net with 26 CONV blocks performance in testing **(upper Right)** and training **(Left)** decreases due to overfitting.

As shown in Figure 10, adding more CONV blocks means increasing both number of parameters and training time significantly. Furthermore, using too many CONV blocks (i.e., Saliency U-Net with 26 CONV blocks) decreased the DSC score due to overfitting.

### 3.6 Exploration of Dilated Saliency U-Net Architecture

In this experiment, we applied different dilation factors in Dilated Saliency U-Net, which captures multi-scale context information on image slices without having to change the number of parameters. As per Figure 9, which visually displays the segmentation results from Saliency U-Net, the boundary delineation is still poor for large WMH regions. Furthermore, we also can see in the same Figure 9 that dilated convolutions help Saliency U-Net to reproduce the shape of WMH regions in more detail. Hence, it is important to know the influence of different dilated convolution configurations in Dilated Saliency U-Net for WMH segmentation.

In order to find the most appropriate dilation factors, we compared different sequences of dilation factors. Figure 3 (c) shows the basic Dilated Saliency U-Net architecture used in this experiment. Only four dilation factors in the encoding part were altered while the rest of the parameters for the training schemes stayed the same. Yu and Koltun suggested to use a fixed filter size for all dilated convolution layers but exponential dilated factors (e.g. 2^0^, 2^1^, 2^2^…) (Yu and Koltun, 2015). Therefore, we assessed “increasing”, “decreasing” and “increasing & decreasing” dilation factor sequences with factor numbers of 1, 2, 2, 4 and fixed filter size of 3 × 3. Details of these configurations are presented in Table 6. From this table, we can appreciate that despite DSU-Net 4221 performed best in DSC score (0.5622), it recorded the lowest sensitivity score. The best sensitivity metric was produced by DSU-Net 1242 (0.4747), but it did not outperform DSU-Net 4221 in DSC score.

**Table 6.**
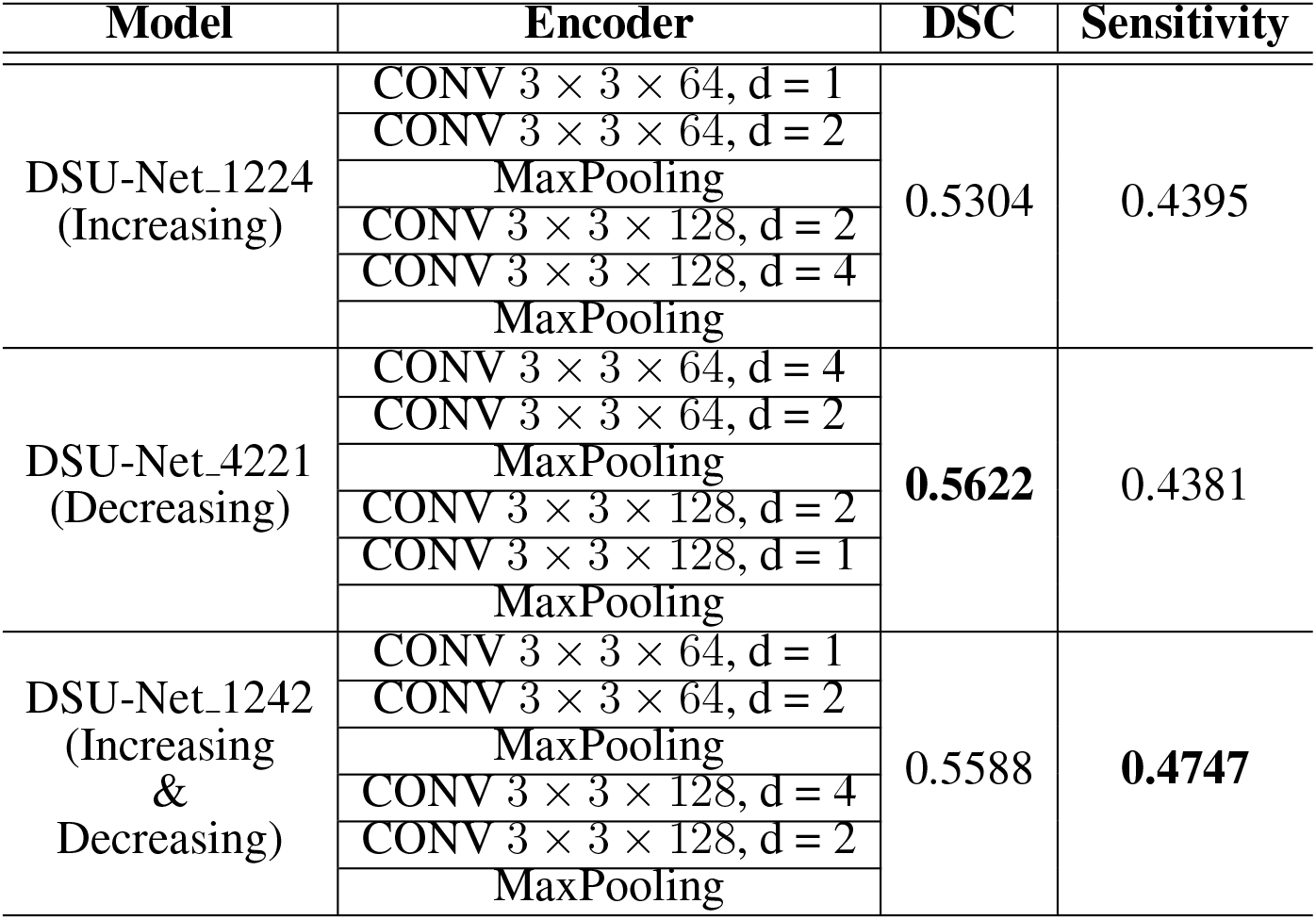
Encoder architecture of Dilated Saliency U-Net with different dilation factors and their performances. Three numbers in the CONV block stans for “filter size × filter size × filter number” and “d” means a dilation factor.

Additionally, we investigated the influence of dilation factors in DSC score performance per WMH volume of MRI scans. Evaluation was conducted on the three groups previously described in Table 3. Figure 11 shows that DSU-Net 1242 outperformed other models in every group. The report of mean, median and standard deviation of DSC score distribution in each group can be seen in Table 4.

**Figure 11.**
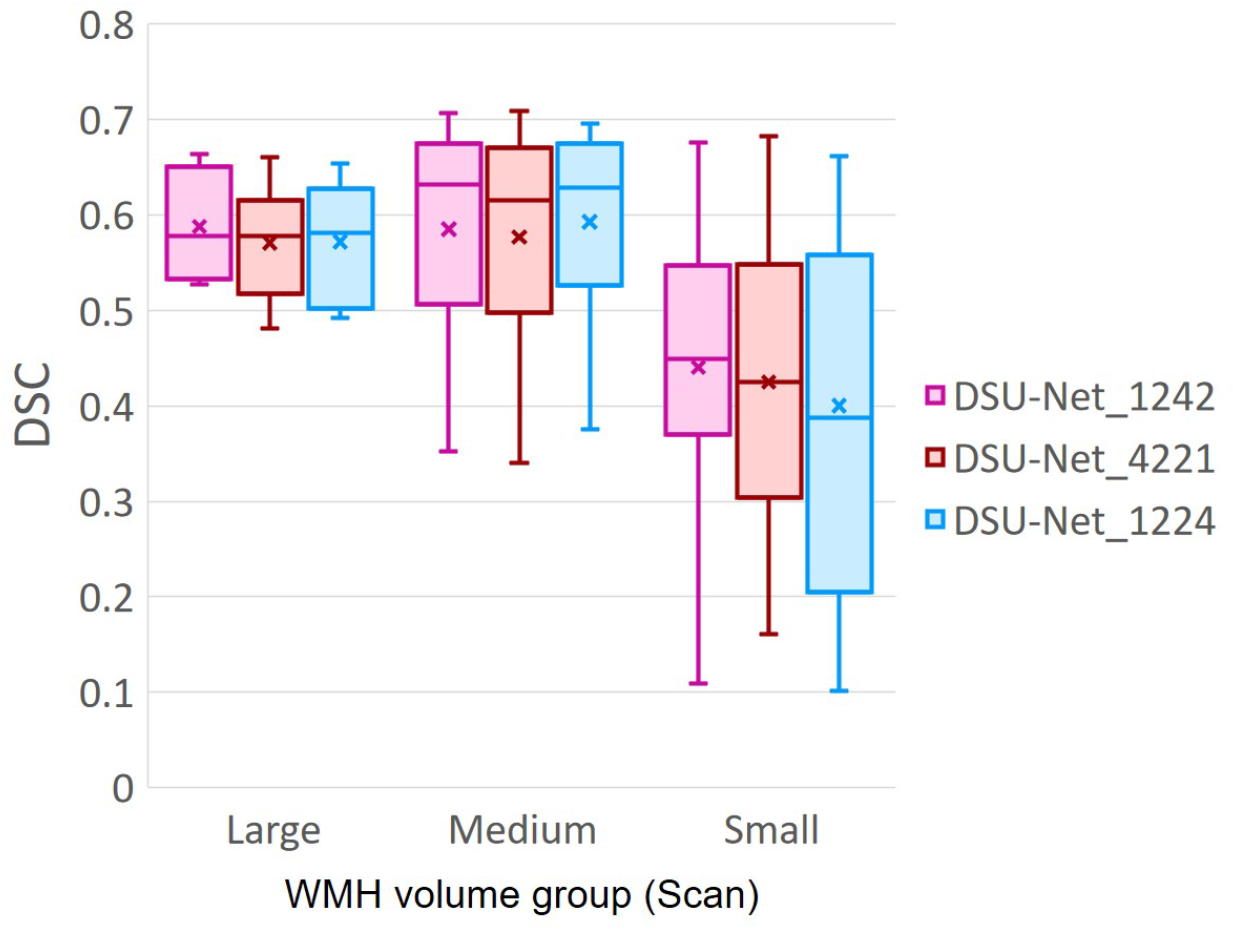
DSC score of three groups based on WMH volume in MRI scans. The group information is described in Table 3. “x” and bar at the middle of box indicate mean and median each. Bottom and top of each box means the first and third quartile.

## 4 DISCUSSION

In this study, we explored the use of IAM as an auxiliary data to train deep neural networks for WMH segmentation. IAM produces a probability map of each voxel to be considered a textural irregularity compared to other voxels considered “normal” (Rachmadi et al., 2019). While incorporating IAM as an auxiliary input data, we compared three deep neural network architectures to find the best architecture for the task, namely U-Net, Saliency U-Net and Dilated Saliency U-Net. It has been suggested that Saliency U-Net is adequate to learn medical image segmentation task with both a raw image and a pre-segmented regional map (Karargyros and Syeda-Mahmood, 2018). The original U-Net did not improve DSC score despite using both T2-FLAIR and IAM as input, but the DSC score from Saliency U-Net was superior to that from the original U-Net trained only with T2-FLAIR. This is because Saliency U-Net is able to learn the joint encoding of two different distributions: i.e. from T2-FLAIR and IAM. Saliency U-Net generated better results than U-Net despite having less parameters. We also found that Saliency U-Net had lower false positive rate compared to U-Net.

Dilated convolution can learn spatially multi-scale context by expanding the receptive field without increasing the number of parameters. We added dilation factors to the convolution layers in the encoding block of Saliency U-Net to improve WMH segmentation, especially due to the high variability in the WMH size. This new model is named “Dilated Saliency U-Net”. Dilated convolution improved both DSC score and sensitivity with shorter training time. Dilated Saliency U-Net also yielded more accurate results in the presence of large WMH volumes and worked well in Medium and Small WMH volume MRI data groups which are more challenging. We identified that dilated convolution is effective when dilation factors are increased and decreased sequentially.

To our knowledge, this is the first attempt of successfully combining dilation, saliency and U-Net. We could reduce the complexity of a deep neural network architecture while increasing its performance through the integrated techniques and the use of IAM. Due to the trade-off between performance and training time, which is proportional to the model complexity, it is crucial to develop less complex convolutional neural network (CNN) architectures without decreasing their performance.

Anomaly detection in the medical imaging field has been broadly studied (Quellec et al., 2016; Schlegl et al., 2017). One of its difficulties relies on the inconsistent shape and intensity of these anomalies. IAM helped the CNN scheme to overcome this problem by providing the localisation and morphological information of irregular regions. We believe it is possible to generate IAM from different modalities of medical images. Thus, the application of IAM is highly expandable to detect different imaging bio-markers involving abnormal intensity values in other diseases.

## DATA AVAILABILITY STATEMENT

The code that corresponds with the experiments described and analysed in this manuscript can be found in https://github.com/hanyangii/DilatedSaliencyUNet.

## CONFLICT OF INTEREST STATEMENT

All authors (Y.J., M.F.R., M.C.V.H., and T.K.) declare that the research was conducted in the absence of any commercial or financial relationships that could be construed as a potential conflict of interest. The funders had no role in study design, data collection and analysis, decision to publish, or preparation of the manuscript.

## AUTHOR CONTRIBUTIONS

Y.J., M.F.R., M.C.V.H., and T.K. conceived and presented the idea. Y.J. and M.F.R. planned the experiments. Y.J. carried out the experiments. All authors provided critical feedback and analysis, and edited the manuscript.

## FUNDING AND ACKNOWLEDGEMENTS

Funds from the Indonesia Endowment Fund for Education (LPDP) of Ministry of Finance, Republic of Indonesia (M.F.R.) and Row Fogo Charitable Trust (Grant No. BRO-D.FID3668413)(M.C.V.H.) are gratefully acknowledged. This project is partially funded by the UK Biotechnology and Biological Sciences Research Council (BBSRC) through the International Partnership Award BB/P025315/1 to M.C.V.H. This study uses data from the Alzheimer’s Disease Neuroimaging Initiative (ADNI) (National Institutes of Health Grant U01 AG024904) and DOD ADNI (Department of Defense award number W81XWH-12-2-0012). ADNI is funded by the National Institute on Aging, the National Institute of Biomedical Imaging and Bioengineering, and through generous contributions from the following: AbbVie, Alzheimer’s Association; Alzheimer’s Drug Discovery Foundation; Araclon Biotech; BioClinica, Inc.; Biogen; Bristol-Myers Squibb Company; CereSpir, Inc.; Cogstate; Eisai Inc.; Elan Pharmaceuticals, Inc.; Eli Lilly and Company; EuroImmun; F. Hoffmann-La Roche Ltd and its affiliated company Genentech, Inc.; Fujirebio; GE Healthcare; IXICO Ltd.; Janssen Alzheimer Immunotherapy Research and Development, LLC.; Johnson and Johnson Pharmaceutical Research and Development LLC.; Lumosity; Lundbeck; Merck and Co., Inc.; Meso Scale Diagnostics, LLC.; NeuroRx Research; Neurotrack Technologies; Novartis Pharmaceuticals Corporation; Pfizer Inc.; Piramal Imaging; Servier; Takeda Pharmaceutical Company; and Transition Therapeutics. The Canadian Institutes of Health Research is providing funds to support ADNI clinical sites in Canada. Private sector contributions are facilitated by the Foundation for the National Institutes of Health (www.fnih.org). The grantee organization is the Northern California Institute for Research and Education, and the study is coordinated by the Alzheimer’s Therapeutic Research Institute at the University of Southern California. ADNI data are disseminated by the Laboratory for Neuro Imaging at the University of Southern California.

## Footnotes

1 http://adni.loni.usc.edu/

2 https://datashare.is.ed.ac.uk/handle/10283/2214

3 https://github.com/febrianrachmadi/lots-iam-gpu

